# All-Flesh Tomato Regulated by Reduced Dosage of *AFF* Provides New Insights into Berry Fruit Evolution

**DOI:** 10.1101/2021.01.04.425284

**Authors:** Lei Liu, Kang Zhang, JinRui Bai, Jinghua Lu, Xiaoxiao Lu, Junling Hu, Chunyang Pan, Shumin He, Jiale Yuan, Yiyue Zhang, Min Zhang, Yanmei Guo, Xiaoxuan Wang, Zejun Huang, Yongchen Du, Feng Cheng, Junming Li

## Abstract

The formation of locule gel is not only an important developmental process in tomato but also a typical characteristic of berry fruit. In this study, we collected a tomato material that produces all-flesh fruits (*AFF*), whose locule tissue remains in a solid state during fruit development. We built genetic populations to fine map the causal gene of AFF trait, investigate the function of *AFF* gene, and identified it as the causal locus conferring the locule gel formation. We determined the causal mutation as a 416-bp deletion that occurred in the promoter region of *AFF*, which reduces the expression dosage of *AFF*. The 416-bp deleted sequence has a high level of conservation among closely related Solanaceae species, as well as in the tomato population. The activity of the 416-bp deletion in down-regulating gene expression was further verified by the relative activity in a luciferase experiment. Furthermore, with the BC6 NIL materials, we reveal that the reduced expression dosage of *AFF* does not impact the normal development of seeds, while produces non-liquefied locule tissue, which is distinct from that of the normal tomatoes in terms of metabolic components. Based on these findings, we propose that the *AFF* gene is the core node in locule tissue liquefaction, whose function cannot be compensated by its paralogs *TAG1, TAGL1*, or *TAGL11*. Our findings provide clues to investigate fruit type differentiation among Solanaceae crops, and also contributes to the breeding application of all flesh fruit tomatoes for the tomato processing industry.

**One Sentence Summary:** The sequence deletion that occurred in the cis-regulatory region of *AFF*—the core node of locule tissue liquefaction determined here—reduced its expression dosage, and produced all flesh fruit tomato.

## INTRODUCTION

Locule gel is a typical characteristic of berry fruit. As the model plant for study of the fruit development and ripening, tomato (*Solanum lycopersicum*) fruit has clear tissue distribution and structure (Czerednik et al., 2015; Huber and Lee, 1986; Joubès et al., 1999). A mass of information on tomato fruit development has been documented but is mainly focused on fruit type, fruit weight, and fruit ripening. There is very limited evidence regarding the regulation of locule gel formation and development (Lamia et al., 2015; Lin et al., 2014; Seymour et al., 2008; Zhu et al., 2018).

Tomato locule tissue, which is the second most abundant tissue in tomato fruit, represents 23% (w/w) of the fruit fresh weight (Mounet et al., 2009). The formation of locule tissue has been proved as a complex process involving a series of physiological and biochemical changes that play a critical role in fruit growth and maturation (Lamia et al., 2015; Lemaire-Chamley et al., 2005; Mounet et al., 2009). Generally, tomato locule tissue derives from the placenta and grows up around the ovules (Davies and Cocking, 1965; Sicard et al., 2010), encloses the developing seeds, undergoes extensive processes of expansion and liquefaction, and turns into a jelly-like homogenous tissue that is composed of thin-walled and giant cells (Atherton and Rudich, 1986; Cheng and Huber, 1996; Joubès et al., 1999). However, the detailed process of its differentiation and formation during the development of tomato fruits remains unknown. The naturally mutated ‘All-flesh fruit’ (briefly named as AFF) tomato does not produce locule gel (jelly-like tissue) surrounding the seeds and completely changes the structure of locule tissue in tomato fruits (Macua et al., 2015; Silvestri, 2006). This might provide an ideal material to uncover the complex mechanism involved in the process of locule development. This mutation also offers several advantages for the tomato processing industry such as its high solid content, improved firmness, long shelf-life, color, and flavor over wild-type tomato (Macua et al., 2015; Silvestri, 2006; Zhang et al., 2019). Therefore, the exploration of AFF may be quite important, not only for the elucidation of berry fruit formation, but also for breeding programs.

Commonly, phytohormones and cell wall-modifying enzymes have been considered to play important roles in locule gel formation. The evidence clearly indicates that IAA, GA, and ABA presented high levels in seeds were transported to the surrounding tissues and then participated in inducing and regulating the development of locule tissue (Kumar and Khurana, 2014; Lemaire-Chamley et al., 2005; Mounet et al., 2009; Sofia et al., 2007). However, it was verified that ethylene and IAA do not control the determination and liquefaction of locule gel in tomato fruit (Brecht, 1987; Gillaspy et al., 1993; Qin et al., 2012). Rather, the formation of locule gel might be related to the ripening and softening processes of fruits because they are accompanied by the dissolution of pectin deglycosylation and hemicellulose, which are the main components of the cell wall matrix, mainly catalyzed by polygalacturonase (PG) and pectinmethylesterases (PME or PE) (Bapat et al., 2010; Cheng and Huber, 1997; Nunan et al., 1998). However, PG and PME mainly change the texture of fruit and do not determine the process of locule gel formation (Tieman et al., 1992; Uluisik et al., 2016). Hence, the initial period of locule gel determination may involve a mechanism that is different from the classic phytohormones or PME – D-galacturonanase scenario.

The well-known floral ‘ABCDE’ model was established to elucidate the molecular mechanism of floral organ development and differentiation. In this model, ABC-class genes are mainly responsible for the formation of sepals, petals, and stamens, while the D-class genes, which belong to the AGAMOUS (AG) family, manipulate the floral organ identity specification, tissue expansion, and fruit maturation in fleshy fruits (Dreni and Kater, 2014; Huang et al., 2017; Huang et al., 2017; Itkin et al., 2010; Vrebalov et al., 2009; Xu and Chong, 2015). Among D-class genes, *AGL1* and *AGL11* specifically contribute to the formation of seeds, the ovule, and funiculus, regulate the expansion and maturation of the carpel and fruit, and promote the development of seeds. For example, as the first set of D-class MADS-box genes reported in petunia, *FLORAL BINDING PROTEIN 7* (*FBP7*) and *FBP11* are expressed specifically in ovule differentiation and formation and also participate in seed and coat development (Angenent et al., 1995; Colombo et al., 1995). Another orthologous gene, *SEEDSTICK* (*STK*; previously *AGL11*), isolated from Arabidopsis, also participates in the initiation and differentiation of ovules and affects seed germination (Ezquer et al., 2016; Favaro et al., 2003; Pinyopich et al., 2003). Suppression of the *STK* orthologous gene *AGL11* triggers seedless fruit in tomato and grape (Ocarez and Mejaí, 2016), whereas overexpression of the tomato *AGL11* gene results in dramatic modifications of flower and fruit organization (Huang et al., 2017). In addition, genes *SHATTERPROOF1* (*SHP1*) and *SHP2* act redundantly with *STK* in promoting ovule identity, control the dehiscence zone differentiation and promote the lignification of adjacent cells at the carpel/ovule boundary (Liljegren et al., 2000; Pinyopich et al., 2003). Similarly, in tomato fruit, *Tomato AGAMOUS-LIKE1* (*TAGL1*), an *SHP* orthologous gene, controls fruit expansion and fleshiness (Vrebalov et al., 2009). More recently, the D-class gene AGAMOUS MADS-box protein 3 (*SlMBP3*)—a paralog of *TAGL1*—showed direct evidence for the liquefaction of tomato locule tissue (Zhang et al. 2019). Moreover, although previous studies indicated that *AGL* genes can control the formation of fleshy fruit, and *slMBP3* impacts the formation of locule gel in tomato, the genetic evolution and regulatory mechanism of the transformation from juiceless to juicy fruit are still unclear.

Moreover, as a common feature, all these D-class genes participate in seed development, such as the *stk* mutant, which reduces seed germination efficiency; *AGL11*, which prevents plants from producing seeds; and *slmbp3* RNAi plants, which develop seeds that are not able to germinate. In contrast, the naturally mutated *aff* as aforementioned produces normal seeds with a high germination rate. Therefore, this *AFF* mutation without gel formation provides additional information for addressing ovule development, especially the locule gel formation, without negative effects on the normal development of seeds.

In this study, the *aff* gene was identified by combining a genetic analysis and map-based cloning approach. We found that a novel structural variant (SV)—a 416-bp sequence deletion—occurred in the conserved cis-regulatory region of the *aff* gene. This deletion led to the suppressed expression of the *aff* gene, and its reduced dosage then caused the all-flesh fruit phenotype but with well-developed seeds. To understand the regulatory pathways and effects on fruit quality caused by the *aff* gene, combined transcriptome and metabolome analyses were performed with the near-isogenic lines (NILs) of the all-flesh tomato material. The metabolic components showed a distinct difference between the mutation and the wild type. Additionally, comparative evolutionary analysis of the *aff* gene and its cis-regulatory sequence in the nightshade family and the tomato population strongly suggests that the *aff* gene is important for fruit development. These findings not only provide useful information for tomato breeding programs but also provide novel insights into the locule gel formation of berry fruits.

## RESULTS

### The All-Flesh Fruit Trait is Controlled by a Single Recessive Locus

To investigate the genetic characteristics of the *aff* genotype, we built an F_2_ population using the *aff* genotype 06-790 as the P1 parent and WT LA4069 as the P2 parent. A fruit trait survey of the F_2_ population showed that the ratio of WT fruit samples to *aff* samples was 150:41, consistent with a 3:1 segregation ratio (chi-squared test: χ2 = 1.176, non-significant) (**Table 1**), which suggests a single recessive genetic model of the all-flesh fruit trait. To further confirm this result, we built a BC_1_P1 population; a survey of the progenies showed that the ratio of WT fruit samples to *aff* samples was 46:42, conforming to a 1:1 segregation ratio (chi-squared test: χ2 = 0.102, non-significant) (**Table 1**). We built a BC_1_P2 population, and all progenies of this population presented WT fruits. In conclusion, these data together confirm that the all-flesh fruit trait is conferred by a single recessive mutation.

**Table 1.**
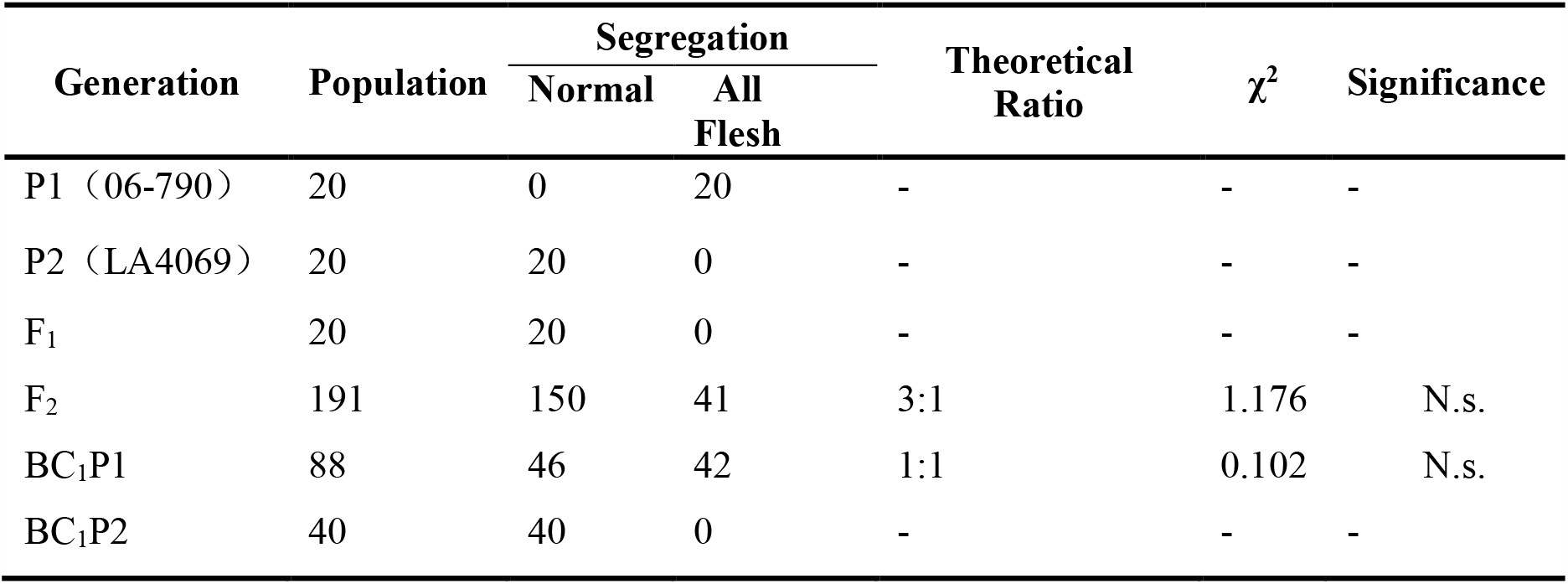
The Traits of the Mature Fruit Locule Tissue of the Populations.

### The Locule cell in *aff* Maintains a Complete Structure during Fruit Ripening

To understand the possible time-point for the locule gel formation, we observed the difference in locule tissue between the *aff* genotype and WT fruits by crosscutting the fruits at an interval of every five days. We found that there was no jelly-like tissue formed in the locule cavity area during the whole development process (fruit setting to ripening) of the *aff* genotype (**Fig. 1a**). In contrast, obvious jelly-like tissue of the WT was observed 25 days after flowering (DAF) and reached complete liquefaction after the mature green (MG) stage (**Fig. 1a**). This jelly-like tissue was composed of distinctly shaped, thin-walled, and highly vacuolated cells. These findings indicate that 25 DAF might be an important time-point for the formation and development of locule gel in tomatoes. We further checked the 25 DAF samples of the WT and all-flesh tomato fruits with paraffin sectioning by microscopic examination. The individual locule cells of the WT tomato fruit continued to collapse and showed a tendency to fracture inter-cellularly within the plane of the cell wall at the MG stage (**Fig. 1b**), which is consistent with previous reports (Cheng and Huber, 1996; Lemaire-Chamley et al., 2005). However, these distinct changes did not occur in the cells of the *aff* locule, while it still maintained a complete structure in *aff* fruits. This made the morphology of the locule tissue in *aff* tomatoes more like that of the placenta tissue. Except for the lack of locule gel formation, the *aff* genotype had a similar ripening process as that of WT tomato.

**Figure 1.**
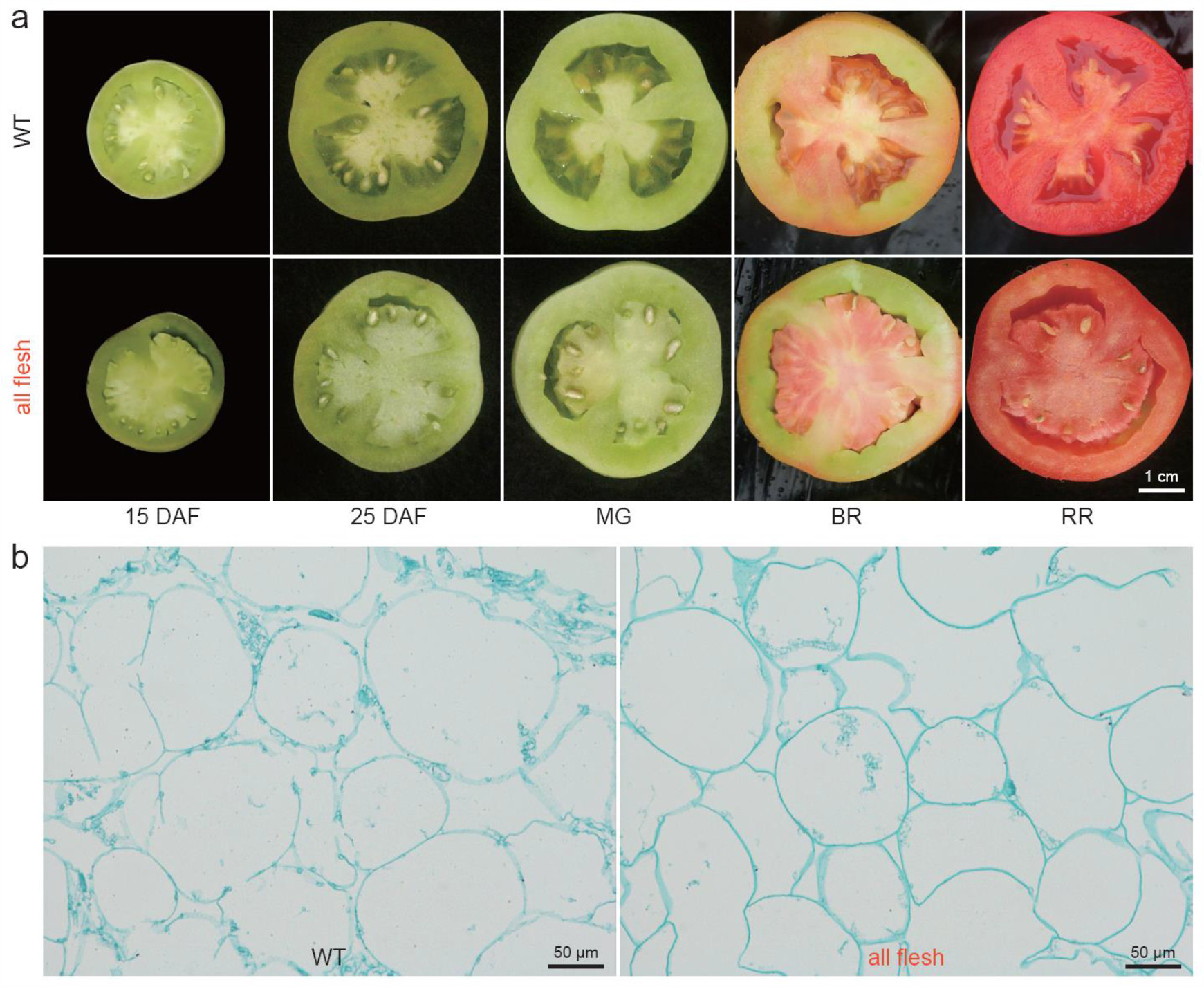
Morphology and Micrograph of the Locule Tissues of Normal (WT) and All-Flesh Fruit Tomato. (**a**) The appearance of locule tissue at different developmental stages of WT tomato LA4069 and all-flesh fruit tomato 06-790. (**b**) The cell structure of locule tissues of WT and all-flesh fruit tomato at their mature green stage. Scale bars: (a), 1 cm; (b), 50 μm.

### A Large Sequence Deletion in the Promoter of the *AFF* gene is identified in All-Flesh Fruit Tomato

A bulked segregant analysis sequencing (BSA-seq) strategy was applied to locate the causal gene in the AFF tomato genome. Using the genome sequence of *Solanum lycopsicum* (SL4.0 ITAG4.0) as the reference, we called out 298,942 SNP variants that were polymorphic between P1 and P2 but homozygous in each of the two parental genomes. These SNP loci were further used in the SNP index analysis (Takagi et al., 2013) with their genotype data from the *aff* genotype pool and the WT pool. Using the measure Δ(SNP index), we detected a significant signal (Δindex=1.448, above the 99% confidence level) located between 37.25 Mb and 37.75 Mb on chromosome six (**Fig. 2a**), and 21 SNPs were located in this region. The average SNP index of the *aff* genotype pool in this region was 0.98, of which 16 SNPs indexes reached 1; the average SNP index of the WT pool was 0.32. The difference in the average SNP index between the two pools was 0.66.

**Figure 2.**
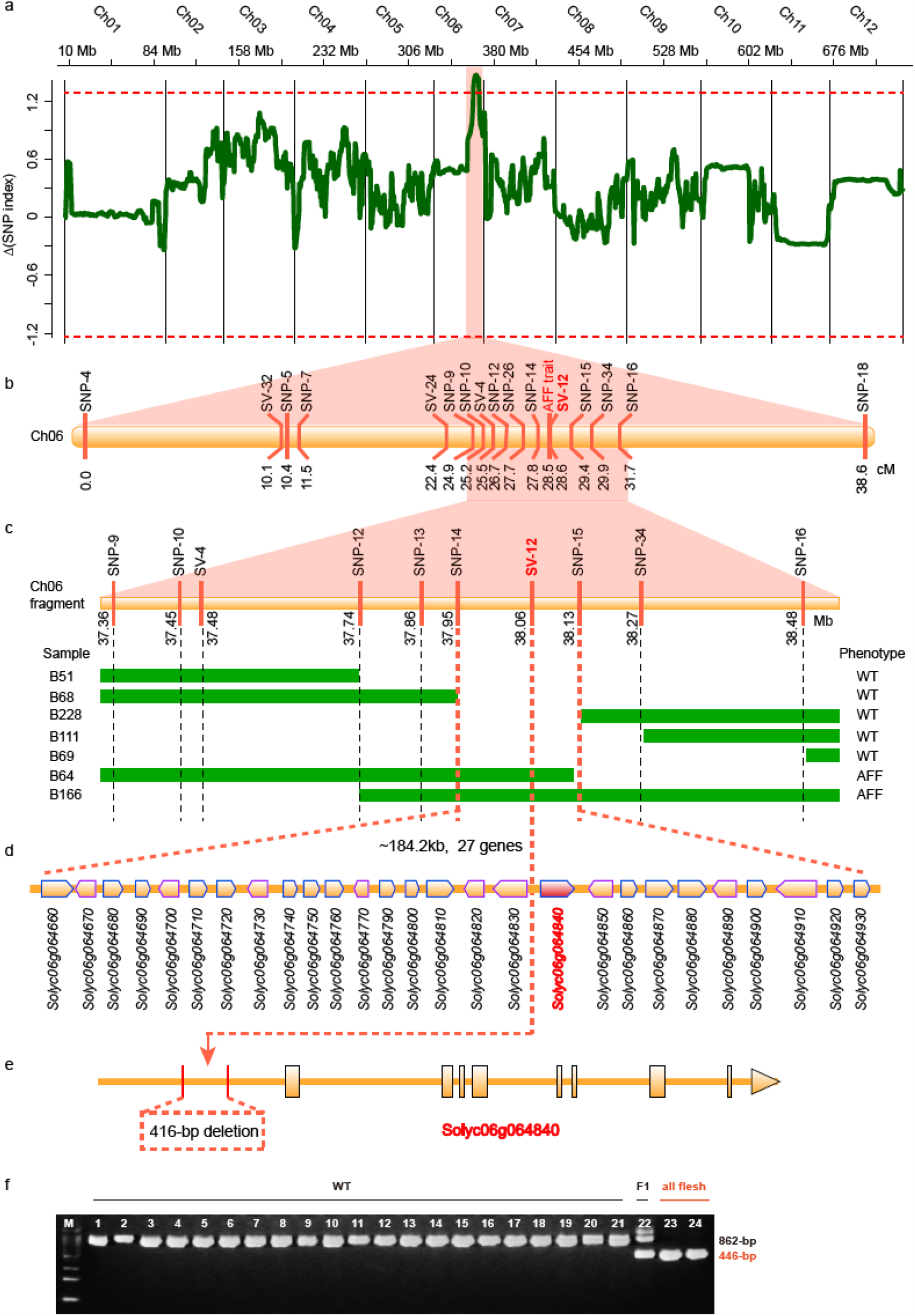
Map-based Cloning of the AFF Gene. (**a**) Δ(SNP index) from BSA-Seq. The *x*-axis is the physical position of tomato chromosomes; the *y*-axis is the value of the SNP-index. (**b**) Initial mapping of the AFF gene using 215 F_2_ plants derived from a cross between 06-790 and LA4069. **(c)** Genotypes and phenotypes of homozygous recombinant plants derived from 249 BC2S1 plants generated by continued backcrossing of 06-790 to H1706 (B51, B68, B228, B111, and B69 are normal lines; B64 and B166 are all-flesh fruit lines). (**d**) Annotated gene models in Tomato SL4.0 ITAG4.0 (H1706) in the mapping region. These local genes are indicated by rectangles with arrows. (**e**) Gene structure of *AFF*. The gray dashed-box represents the SV of 416-bp deletions in the cis-regulatory region of the *AFF* gene. (**f**) The PCR results of different tomato varieties or lines using the marker SV-12 designed by the 416-bp deletion. M: 100-bp DNA ladder.

Genetic linkage analysis of two populations (F_2_ with 215 individuals and BC_2_S_1_ with 249 individuals as listed in the methods) was used to fine-map the *AFF* gene. Molecular markers (**Table S1**) were selected from these polymorphic SNPs and SVs between P1 and P2 and were genotyped by PCR and KASP. The results showed that the *AFF* gene was mapped to the same region as that in BSA-Seq (**Fig. 2b)**. Based on the physical location of these markers, as well as the genotype of each individual derived from the BC_2_S_1_ population, the *AFF* gene was finally mapped between markers SNP-14 and SNP-15, corresponding to the physical location of SL4.0ch06:37,945,500-38,129,705, which is about 184.2 kb and harbors 27 genes (**Fig. 2c and 2d**). The variants in this region were further deciphered. Only a 416-bp deletion was found in SL4.0ch06:38,062,128-38,062,543, lying in the intergenic region. Following the gene model information in the tomato reference genome, we found that this 416-bp deletion was located 1,775 bp upstream of the gene *Solyc06g064840* (**Fig. 2e**). Based on the 416-bp deletion, a marker named SV-12 was designed, and two populations were screened by this marker. We found that SV-12 completely co-segregated with all-flesh individuals (**Fig. 2f**). This evidence determined *Solyc06g064840* as the candidate gene, which belongs to the AGAMOUS-like MADS-box transcription factor gene family and is also named *SlMBP*3. *SlMBP3* is specifically expressed in the developing locule (include the seeds) of tomato fruits (**Fig. S1-2**) (Fernandez-Pozo et al., 2017; Koenig et al., 2013).

### Gene Editing of *AFF* Confirmed its Function in locule Gel Formation of Tomato

To prove that *Solyc06g064840* is the causal gene of the *aff* tomato genotype, an 18-bp sgRNA that targeted the second exon of the *AFF* gene was designed to construct a CRISPR-Cas9 expression vector MSG8124/8125 and then introduced into the *S. lycopersicum* cv. Micro-tom (wild type) by *Agrobacterium tumefaciens* (**Fig. 3a-b**). These transgenic plants were confirmed by PCR amplification and DNA sequencing (**Table S2**). As shown in **Fig. 3b**, the *AFF*-Cr5 mutant produced the expected all-flesh fruits without the jelly-like substance compared to the WT. We also constructed the recombinant plasmid 35S::5’UTR+ aff-CDs::GFP and introduced it into Micro-tom to obtain transgenic plants with over-expression of the *Solyc06g064840* gene. As shown in **Fig. 3c**, the *AFF*-overexpression transgenic T_1_ homozygous lines developed larger jelly-like substances in the locule compared to the normal locule gel in the WT Micro-tom tomato fruits. These results indicate that the *AFF* gene does possess the key function in locule gel formation of tomato fruits.

**Figure 3.**
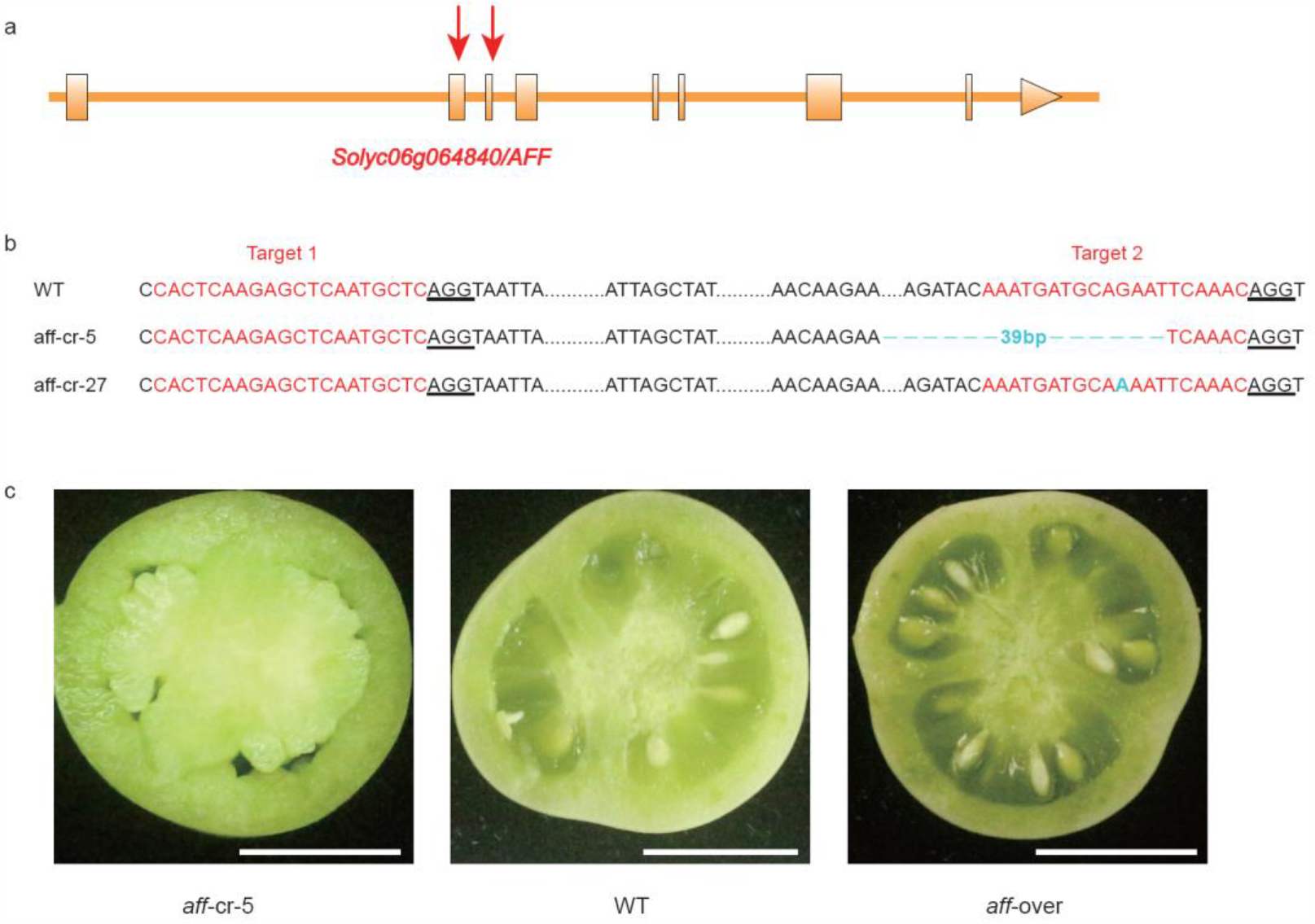
Characterization of CRISPR/Cas9-*aff* (*aff*-cr) Lines and Over-expression (*aff*-over) Lines. (**a**) Schematic illustrating single-guide RNA targeting the *AFF* coding sequence (red triangle). **(b**) *aff*-cr mutants generated using CRISPR/Cas9. The red lines indicate the target sites of the guide RNAs. The nucleotides underlined in black bold font represent the protospacer-adjacent motif (PAM) sequences. *aff*-cr alleles identified by cloning and sequencing PCR products of the *AFF-*targeted region from two T_0_ plants under the MicoTom background. (**c**) Representative fruit transection from CRISPR/Cas9-*aff* (*aff*-cr) lines compared with the wild-type (WT) and over-expression (aff-Over) lines at 25 DAF. Scale bars: 1 cm.

### The Deleted Promoter Sequence Shows Strong Conservation

To understand the detailed function of the 416-bp deletion, the 2 kb sequence (416-bp deletion included) was analyzed by the promoter prediction tool TSSP of the PlantProm DB database (Shahmuradov et al., 2012) and the PlantCARE database (Lescot et al., 2002). We found that functional elements including the TATA box and CAAT box (**Table S3**) were involved in this deleted sequence. Therefore, it is interesting to investigate the conservation status of the 416-bp sequence across Solanaceae crops. To do this, we selected five genomes from four Solanaceae crops, including two tomato genomes *S. lycopersicum* and *S. pennellii*, as well as genomes of potato (*Solanum tuberosum*), capsicum (*Capsicum annuum*), and eggplant (*Solanum melongena*). We then determined the syntenic orthologous genes of *AFF* in the five genomes selected (**Methods**), which are *Sopen06g023350, Sotub06g020180, Capang01g002169*, and *Sme2*.*5_02049*.*1_g00007*.*1*, in *S. pennellii, S. tuberosum, C. annuum*, and *S. melongena*, respectively. The promoter sequences of these five syntenic genes were extracted from corresponding genomes and aligned using the MUSCLE tool (Edgar, 2004). Based on the results of multiple sequence alignment, we estimated the conservation level of these promoter sequences. As shown in **Fig. 4a**, using the revised π as the measure (**Methods**) and the threshold of 0.3, we determined that five main local regions showed a relatively higher conservation level (low mismatch ratio in multiple sequence alignment) in the promoter sequences of these Solanaceae crops. These five regions should have important roles in regulating the expression of associated genes. Moreover, the 416-bp sequence deletion was exactly located at one of the two most conserved regions (the blue bars in **Fig. 4a**). This suggests that the deletion may have a large effect in altering the expression of the gene *AFF* in *aff* tomato.

**Figure 4.**
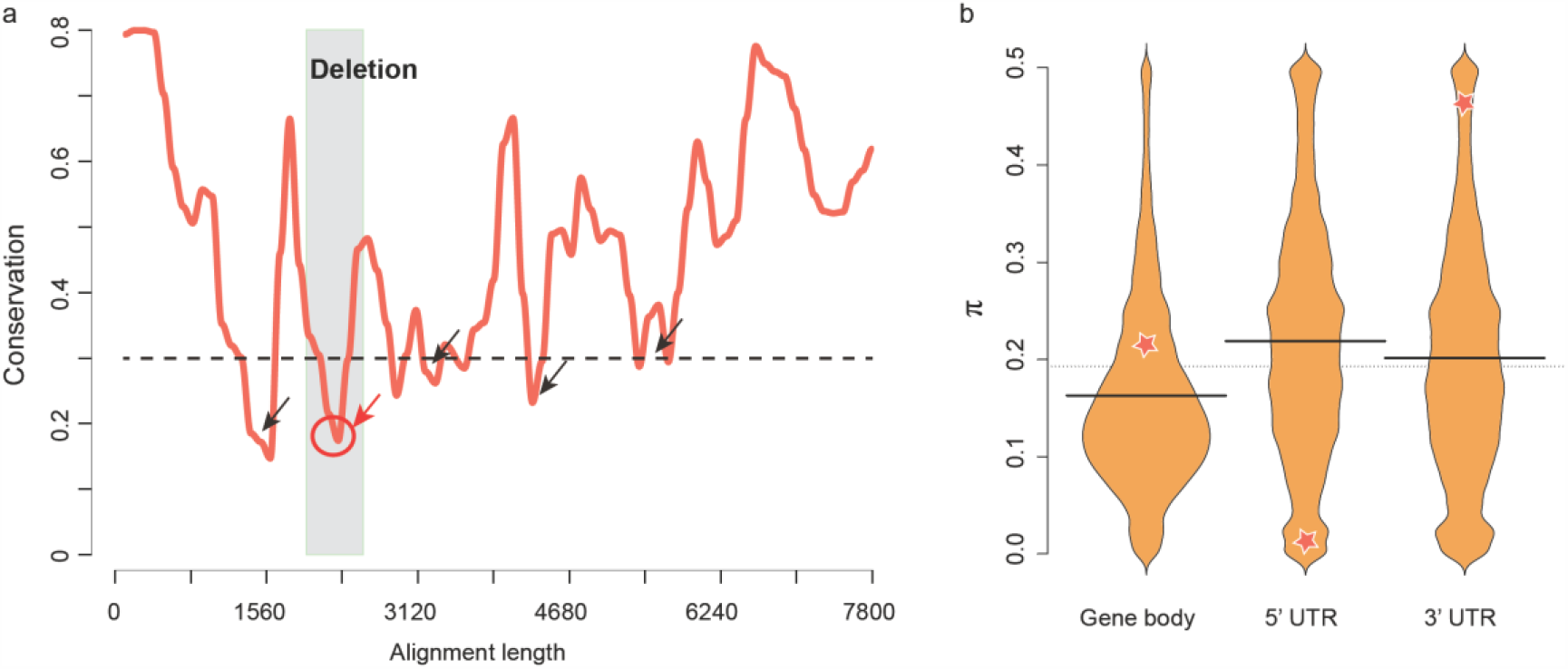
The Sequence Conservation of the *AFF* Promoter in *Solanaceae* Species and the Tomato Population. (**a**) Sequence conservation of *AFF* orthologous genes among five *Solanaceae* species. (**b**) Beanplot of π values for the three regions: the gene body, 5’UTR, and 3’UTR of all genes. The 5‘UTR region of gene *AFF* shows strong conservation compared to other genes in the tomato population. The yellow stars show the π values of the three regions of the *AFF* gene.

To further check the conservation and importance of the promoter region of *AFF* within the tomato population, we analyzed its sequence diversity using the population variome dataset on 360 tomato accessions that was published previously (Lin et al., 2014). Generally, we estimated the sequence diversity (selection sweep) by calculating π for regions of the gene body, 5’UTR, and 3’UTR for each of the 33,562 tomato genes (**Methods**) and checked the selection strength, i.e., the diversity level of the *AFF* gene under the background of all tomato genes. As shown in **Fig. 4b**, the yellow star shows the location of the *AFF* gene in the frequency distribution of the π values calculated from all genes. For *AFF*, the π value of the gene body is 0.21, slightly larger than the mode value of all genes, while its 3’UTR has a π value of 0.46, which indicates higher diversity, whereas its 5’UTR has a π value of 0.026, less than 93.80% of all other genes, suggesting strong selection pressure against mutations in the promoter region of the *AFF* gene in the tomato population. These findings together support the importance of sequence conservation in the promoter region of *AFF*, which further suggests that the 416-bp deletion may have a significant impact on the function of *AFF*.

### The Promoter Sequence Deletion Down-regulates the Expression Level of the *AFF* gene

We investigated the expression of *AFF* by quantitative real-time PCR assay and the dual luciferase reporter system. First, using the marker of the 416-bp deletion variant, we selected two *aff* lines (BA-130 and BA-150) and two WT lines (BA-124 and BA-128) from the BC_2_S_1_ population of P1 (06-790) and P3 (H1706). We then performed qRT-PCR analysis to measure the expression variation of *AFF* in different developmental stages of locule tissues from the four BC_2_S_1_ lines, together with their parental materials P1, P3, and F_1_, as well as another *aff* line 09-1225 and the WT line P2 (LA4069). As shown in **Fig. 5a**, in all of these samples, the gene was highly expressed on seven DAF and ten DAF and significantly decreased on 15 DAF, which was synchronous with the differentiation of locule tissues, and is consistent with previous reports (Fernandez-Pozo et al., 2017; Koenig et al., 2013). More importantly, the expression of *AFF* was significantly lower in samples of *aff* tomatoes than in WT fruit samples. The all-flesh BC_2_S_1_ lines BA-130 and BA-150 had a significantly lower expression of *AFF* than the WT fruit BC_2_S_1_ lines BA-124 and BA-128 (**Fig. 5a**). We further evaluated the transcriptional activity of the promoter sequences of *AFF* in the WT and *aff* fruit samples, i.e., promoter sequences with or without the 416-bp deletion, by experiments of the relative activity of luciferase (the ratio of luc to Rluc). The relative activity of luciferase of the all-flesh fruit promoter was significantly lower than that of the WT promoter (**Fig. 6a)**. All of these results indicate that the 416-bp deletion in the promoter region can significantly decrease the transcription level of *AFF*, and the down-regulated expression of *AFF* then regulates the formation of all-flesh fruit tomatoes.

**Figure 5.**
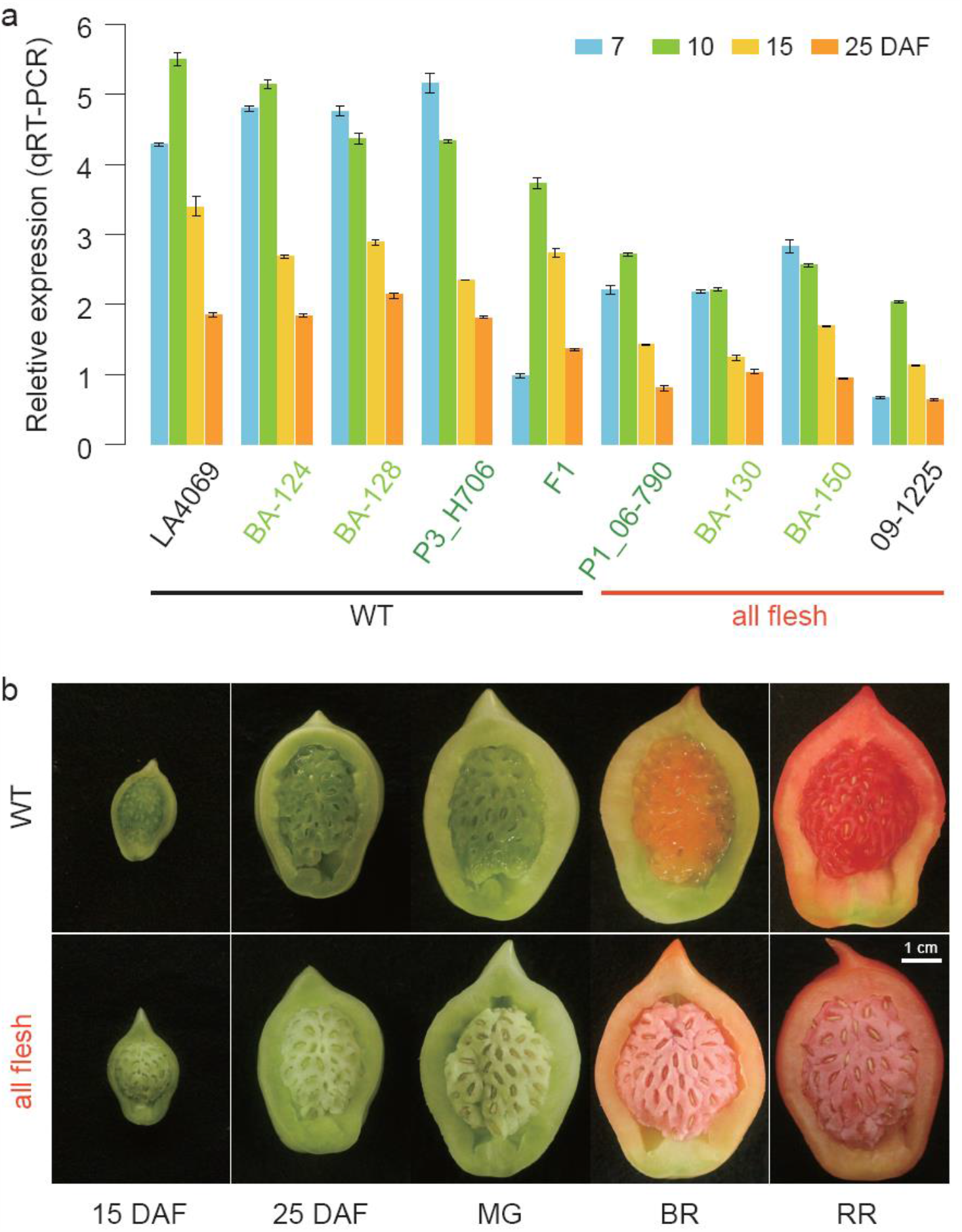
The Expression of Gene *AFF* and the Phenotypes of Locule Tissues of WT and All-Flesh Tomato Fruits at Different Development Stages. (**a**) qRT-PCR of *AFF* transcripts in different locule tissues and stages from 7 to 25 days after flowering (DAF). BA-130 and BA-150, all-flesh lines derived from BC_2_S_1_ plants generated by the continued backcrossing of 06-790 to H1706. 09-1225 and 06-790 are all-flesh cultivars. 06-790×H1706, F_1_ progeny, the all-flesh line 06-790 was crossed to wild-type H1706. H1706 and LA4069 are normal cultivars obtained from TGRC. BA-124 and BA-128 are normal lines derived from BC2S1 plants generated by continued backcrossing of 06-790 to H1706. Note: To normalize the expression data, the *SIFRG27* (*Solyc06g007510*) gene was used as the internal control (Cheng et al., 2017). The bars represent the standard deviation. (**b**) The longitudinal section of fruit locule tissue at different stages of the WT and *aff* NIL (PA-1) created by backcrossing of 06-790 to H1706 for six generations followed by two generations of selfing. Scale bars: 1 cm.

**Figure 6.**
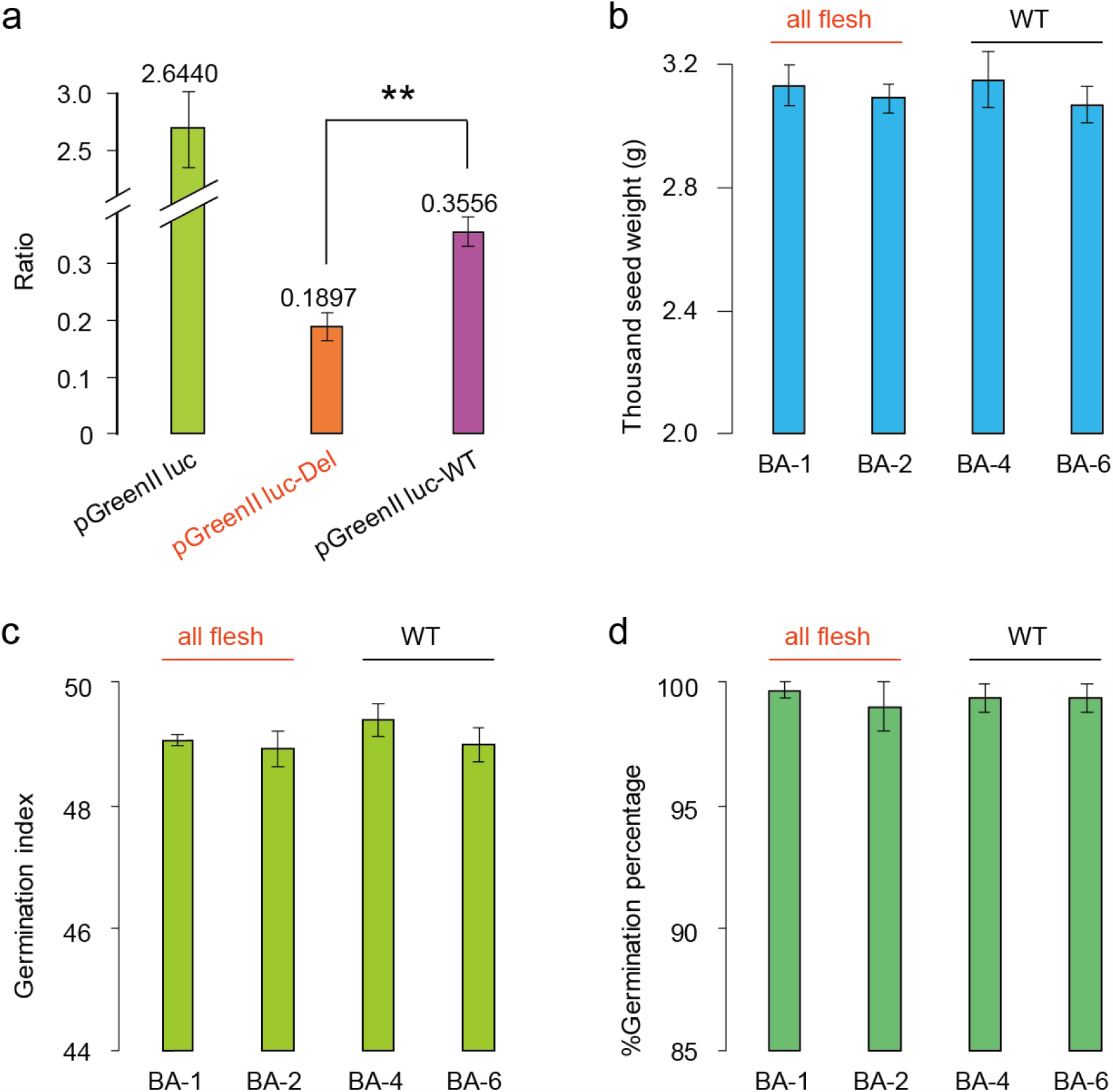
The Ratio of Firefly and Renilla Luciferase Signals, as well as the Thousand-Seed Weight and the Seed Germination of *aff* NILs. (**a**) Relative luciferase activity (the ratio of luc to Rluc) of the two constructs. pGreenll luc, the blank vector with the 35S promoter; pGreenll luc-Del, the vector with the *aff* promoter; pGreenll luc-WT, the vector with the *AFF* promoter. Different letters above the bars indicate statistically significant differences. **: P < 0.01 (Student’s t test). (**b**) The thousand-seed weight of four *aff* NILs. (**c**) The germination index of four *aff* NILs.**(d)** The seed germination percentage of four *aff* NILs. BA1-1 and BA2-1 are all-flesh lines derived from BC_6_S_2_ plants generated by 06-790 continued backcrossing to H1706; BA4-1 and BA6-1 are normal lines derived from BC_6_S_2_ plants generated by 06-790 continued backcrossing to H1706. The bars represent the standard deviation.

Furthermore, we checked the locule tissues of *aff* and WT tomato fruits using the near-isogenic lines (NILs). The NILs were generated by back-crossing the *aff* tomato material 06-790 to H1706 for six generations (**Methods**), assisted by molecular selection of the 416-bp deletion marker SV-12. We surveyed these lines with the homozygous status of the 416-bp deletion and observed that they all produced all-flesh fruit, which is distinct to the WT fruit of H1706 (**Fig. 5b**), in which the locule tissues of all-flesh tomatoes maintain a solid state during fruit development, without any jelly tissue formation. Additionally, we also checked the seed characteristics, thousand-seed weight, and the seed germination activity using the all-flesh fruit tomato NILs; the seed structure or appearance did not differ between the *aff* fruit lines and the WT fruit lines (**Fig. S3**). All the *aff* NILs had complete seed hair and coat. As shown in **Fig. 6b-d**, the all-flesh fruit lines BA-1 and BA-2 had a similar thousand-seed weight, germination index, and germination rate as that of WT fruit lines BA-4 and BA-6. These results suggest that the deletion stops the formation of gel in tomato but does not impact the function of *SlMBP3*/*AFF* involved in the normal development of seeds. The deletion mutation in the all-flesh fruit was different from that of the *aff*-cr5 plants that cannot produce seeds and the *SlMBP3* RNAi plants whose seeds cannot germinate (Zhang et al., 2019).

### The *AFF* Mutation Largely Alters Gene Expression and Metabolic Components

The reduced expression of *AFF* shut down downstream locule tissue liquefaction-related biology pathways. A low dosage of AFF had an impact on systematic gene expression variations in the locule tissue of *aff* tomato, i.e., the expression of more genes was down-regulated. We compared whole genome gene expression patterns between tomato materials HZ106 (WT) and its NIL BA-1 (*aff* line) whose *AFF* gene was replaced by the mutated one with the 416-bp deletion in its promoter region. mRNA-seq analysis were performed on two tissues, the locule and placenta, for both the WT and *aff* line, at three time points, i.e., 10, 15, and 25 DAF. Generally, we found that genes belonging to GO (gene ontology) terms related to lipid metabolism, plant-type cell wall, phyto-hormones, metabolism and catabolism, flavonoid biosynthesis, glucosyltransferase activity, and nutrient reservoir activity, were enriched in these differentially expressed gene sets (**Table S4**), as well as KEGG pathways including sugar metabolism and phyto-hormone biosynthesis (**Table S5**). Among the top 50 enriched GO terms, 1,110 genes were down-regulated, while only 359 genes were up-regulated (**Table S4**). In the KEGG pathway ‘MAPK signaling’, 55 differentially expressed genes were involved, with 42 genes down-regulated and 13 genes up-regulated. These results clearly show a large number of genes whose expression was altered—mostly down-regulated—by the reduced expression of *AFF* in the *aff* tomato fruit line; these suppressed pathways may then prevent the locule tissues from liquefying. Furthermore, we performed detailed comparisons between pair-wise transcriptome datasets. When comparing gene expressions between locule and placenta tissues of the WT tomato (group 1), we found that genes involved in the GO terms lipid transport, apoplast, flavonoid metabolism, transferase, and hydrolase activity or KEGG pathways such as metabolism, protein kinase, and phyto-hormone were enriched (**Fig. S4**). However, the GO terms lipid transport and flavonoid metabolism were not enriched in differentially expressed genes between the locule and placenta tissues of the *aff* tomato lines (group 2). Additionally, there was over-representation of the GO terms phloem or xylem, as well as symporter activity or transmembrane transmission-related genes that were differentially expressed in group 2 (**Fig. S5**). To focus on the differences between the locule tissues from the WT and *aff* tomato lines (group 3), we further compared their differentially expressed genes and found that similar GO terms or KEGG pathways as those observed in group 1 were enriched **(Fig. 7a-b)**. Further, the GO terms DNA replication, plasma membrane, photosystem II, plant-type cell wall, glucosyltransferase activity, as well as nutrient reservoir activity were specifically enriched in group 3 (**Table S6** and **Fig. 7a**). Additionally, we also compared the gene expression difference between the placenta issues from the WT and *aff* tomato lines (group 4). However, the aforementioned KEGG pathways or GO terms were not enriched or differentially expressed in group 4 (**Fig. S6**). These results together suggest that the reduced expression of *AFF* mainly down-regulated the expression of genes involved in DNA replication, phyto-hormone metabolism, photosynthesis, sugar metabolism, and MAPK signaling, which further prevented the locule liquefaction process that normally occurred during the differentiation of locule tissue in the placenta of WT tomato fruits.

**Figure 7.**
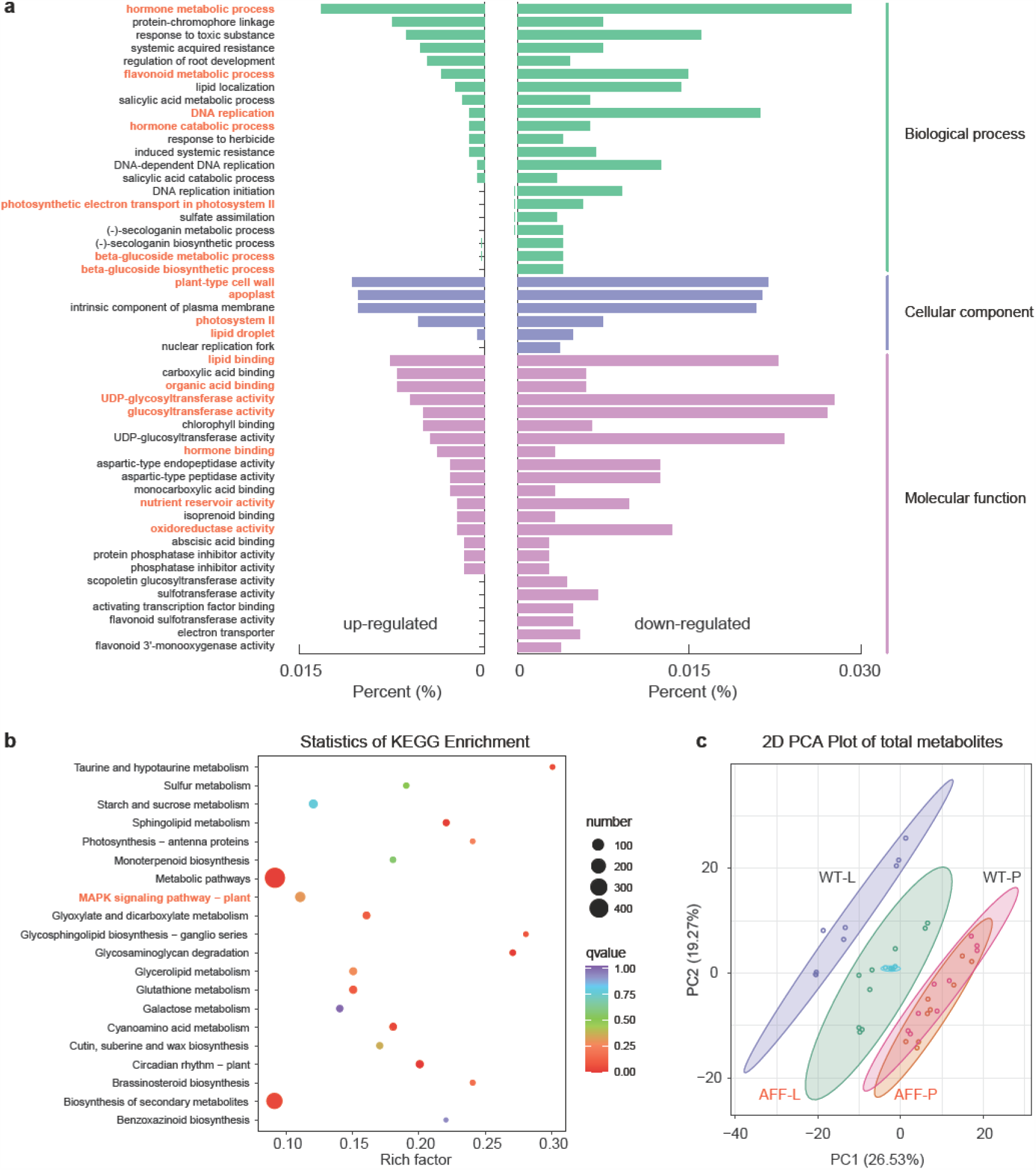
The Differential Gene Expression and Metabolite Contents between Fruits from the All-Flesh and WT Tomatoes. The significantly enriched GO terms (**a**) and KEGG pathways (**b**) of differentially expressed genes between locule tissues from the all-flesh fruit and WT tomatoes. (**c**) The principal component analysis of metabolites from the locule and placenta of WT and all-flesh fruit tomatoes. WT-L: locule tissue of WT tomatoes; WT-P: placenta tissue of WT tomatoes; AFF-L: locule tissue of all-flesh fruit tomatoes; AFF-P: placenta tissue of all-flesh fruit tomatoes.

Genes that have key functions in the liquefaction of locule tissues showed strong expression differentiation between the locule tissues of the *aff* and WT tomato fruits. Genes involved in hydrolases, phyto-hormone metabolism, and DNA replication, were reported to be involved in the regulation of locule tissue liquefaction of tomato fruits (Christian et al., 2014; Huber and Lee, 1986; Mounet et al., 2009; Takizawa et al., 2014; Uluisik et al., 2016). We examined the differentially expressed genes between tissues from the WT and the *aff* line. Among the 188 genes that showed stable and strong differential expression, 122 genes were down-regulated, while 66 genes were up-regulated in *aff* tomatoes in comparison to the WT (**Table S7**). Many of these differentially expressed genes are locule tissue liquefaction process-related genes. First, six gibberellin-related genes were down-regulated in *aff* tomato fruit. Among them, four are gibberellin-regulated proteins, while one is involved in the gibberellin biosynthesis process, and the last one is involved in the gibberellic acid-mediated signaling pathway (**Table S7**). We also found six auxin-related genes that were differentially expressed, with the auxin repressed protein up-regulated, while the other five genes, auxin transporters, auxin responsive protein IAA9, auxin-related genes from the GH3 gene family and those involved in the auxin signaling pathway, were down-regulated in *aff* tomato. These phyto-hormone-related genes should play key roles in the liquefaction of locule tissue in WT tomato. Second, there were three copies of cytochrome P450 genes whose expressions were down-regulated in *aff* tomato fruits (**Table S6**), indicating a low level of energy-related activity in *aff* fruit tomato. Third, pectinesterase (PE) that is strictly regulated and functions in the softening of tomato fruits was largely down-regulated in *aff* tomatoes. This may have an important impact on the solidness of *aff* tomato fruits. Fourth, we identified two copies of xyloglucan endo-transglycosylase (XET)-related genes that were down-regulated in *aff* tomatoes. XET is involved in the induction of fruit ripen and softening, its down-regulation should hinder the softening of *aff* tomato fruits. Fifth, we found three copies of glycosyl hydrolase genes that were down-regulated in *aff* tomato fruit, indicating suppressed metabolism of glycolysis in *aff* tomato compared with the WT tomato. Interestingly, the gene *PAR2* (*Solyc01g008550*), which encodes phenylacetaldehyde reductase, was up-regulated in the locule tissue of *aff* fruit tomato compared with the WT tomato (2.50-, 1.87-, and 6.59-fold changes at 10, 15, and 25 DAF, respectively). Phenylacetaldehyde reductase has been reported as the key gene catalyzing the synthesis of the aroma volatile 2-phenylethanol in tomato; its up-regulated expression in the locule makes *aff* tomato more specific in flavor quality relative to WT tomato (Tieman et al., 2006), thus endowing the *aff* tomato with a flavor advantage for the food processing industry (Macua et al., 2015). Moreover, we found that *TOMATO AGAMOUS 1* (*TAG1, Solyc02g071730*) and *TAGL1* (*Solyc07g055920*)—both are paralogs of *AFF* and show high levels of sequence homology (**Fig. S7**)—were up-regulated in the locule tissue of *aff* tomato compared with the WT tomato, which may suggest that the feedback compensation expression of *TAG1* and *TAGL1* under down-regulated expression of *AFF*. Additionally, *TOMATO AGAMOUS LIKE 11* (*TAGL11, Solyc11g028020*), the paralog with the highest homology to *AFF* in tomato (Huang et al., 2017; Zhang et al., 2019) (**Fig. S7**), showed no expression difference between *aff* and WT tomatoes; thus, *TAGL11* may not be one member of the feedback compensation loop shared by *TAG1, TAGL1*, and *AFF*. However, the fact that the up-regulated expression of *TAG1* and *TAGL1*, as well as the stable expression of *TAGL11*, did not compensate the low dosage of AFF to recover from the all-flesh fruit trait suggested that the mediation of locule tissue liquefaction is unique to the *AFF* gene and not to its three paralogs *TAG1, TAGL*, and *TAGL11*. These tomato fruit development-related genes—whose expressions were largely altered as a subsequent effect of the reduced expression of *AFF*—are the key genes that suppressed the locule tissue liquefaction in *aff* tomatoes.

Our metabolic data support the results observed in the mRNA-seq analysis. We measured the metabolites and their quantities in tomato fruits of the WT and *aff* line (**Methods**). Principal component analysis (PCA) of metabolites from the WT and *aff* line (**Fig. 7c**) showed that the placenta tissues from the *aff* and WT tomato lines had similar metabolic components. However, the metabolites of locule tissue were different from those of the placenta tissue in the WT or *aff* tomato. More importantly, the pattern of locule metabolites in the *aff* tomato was located in between that of the placenta and locule tissues of the WT tomato. This indicates that the down-regulated expression of *AFF* changed the metabolic components in the locule tissue of *aff* tomato. Furthermore, we investigated the metabolites whose contents were changed in the *aff* tomato compared to those in the WT tomato and found higher levels of flavonoids and lipids (**Table S8**) in the *aff* tomato, whereas there were more alkaloids and phenolic acids (**Table S8**) in the WT tomato than in the *aff* tomato. The differences in the metabolic components were caused by the down-regulated expression of *AFF* and downstream large-scale gene expression variations, which further resulted in the distinct fruit quality, as taste and flavor, of the *aff* tomato, compared to the WT tomato.

## DISCUSSION

Locule gel liquefaction is not only a significant process in development and ripening but also a typical characteristic of tomato fruit. In this study, the causal gene *AFF* of the all-flesh fruit trait and the 416-bp key deletion mutation in the cis-regulatory region of *AFF* were determined, and we further found that the liquefication function of locule tissue is mediated uniquely by *AFF*, while the expression dosage of *AFF* is crucial for locule tissue liquefaction. *AFF* belongs to the AGAMOUS gene family and contains typical MADs-box domain; its paralogous genes in tomato are *TAG1, TAGL1*, and *TAGL11*, which have a high sequence homology (**Fig. S7**). These genes and their orthologs were found to play important roles in ovule differentiation and formation, participate in seed and coat development, or regulate the expansion and ripening processes of the carpel and fleshy fruit in many species (Angenent et al., 1995; Colombo et al., 1995; Itkin et al., 2010; Pan et al., 2010; Favaro et al., 2003; Huang et al., 2017; Ocarez and Mejaí, 2016; Vrebalov et al., 2009). Recently, using reverse genetics, Zhang et al. (2019) showed that *AFF* (*slmbp3*) had impacts on locule tissue liquefaction and seed formation in tomato, while the seeds from RNAi plants lost germinability. However, in our *aff* genotype, the 416-bp deletion down-regulated the expression of *AFF*, inhibiting locule gel formation but producing normal seeds.

The *AFF* gene functions as a core node of locule liquefaction whose function cannot be compensated by its paralogs *TAG1, TAGL1*, or *TAGL11*. The cis-regulatory sequence deletion mutation of the *AFF* gene caused the differential expression of many important genes. Among them, we observed that the expressions of *TAG1* and *TAGL1* were significantly up-regulated in *aff* tomato, accompanied by the down-regulated expression of *AFF*. Another paralog, *TAGL11* (**Fig. S7**), which is involved in fleshy tissue differentiation of tomato (Huang et al., 2017), showed stable expression between *aff* and WT tomatoes. This suggests that *AFF, TAG1*, and *TAGL1* may share one expression-feedback loop, without *TAGL11*. More importantly, considering the fact that the up-regulated expression of *TAG1* and *TAGL1* and the stable expression of *TAGL11* did not recover the normal liquefied locule tissue from the all-flesh fruit tomato, the function of mediating locule tissue liquefaction should be unique to the gene *AFF*, but not its paralogs, i.e., the other D-class genes in tomato (**Fig. S7**). Furthermore, based on metabolomics analysis, we found that the pattern of locule metabolites in the *aff* tomato was located in between that of the placenta and locule tissues of the WT tomato, which indicates that tomato locule tissue is derived from the placenta, which is formed from the development of the carpel (Davies and Cocking, 1965; Lemaire-Chamley et al., 2005; Pedro et al., 1991; Sicard et al., 2010). The process is regulated by D-class genes in the classical ‘ABCDE’ flower development model and is consistent with the locule gel formats along with the degradation of the cell wall matrix (Brecht, 1987; Chevalier et al., 2011; Joubès et al., 199 9; Lemaire-Chamley et al., 2005).

The reduced dosage of the AFF protein caused by a 416-bp cis-regulatory deletion is the key factor that promoted the formation of the all-flesh fruit trait. The dosage of gene expression has been proved to play an important role in the variation of plant traits, especially for the floral organ identity that determines genes. Up- or down-regulation of the expression of one ABCDE-class gene may easily shift the boundaries between different types of floral organs (Ito et al., 2007; Wang et al., 2016; Wuest et al., 2012). For example, a dosage imbalance between B- and C-class proteins can change stamen morphology (Liu et al., 2018), while the expression variation of *TAGL1* and *TAGL11* can also affect the development of tomato seeds and the fleshy characteristic (Gimenez et al., 2016; Huang et al., 2017; Ocarez and Mejaí, 2016; Vrebalov et al., 2009). Besides floral-determining genes, another example shows that gene editing of different loci in the promoter region of tomato genes resulted in fruits with different sizes (Rodriguez-Leal et al., 2017). The structural variant (SV) has been found as one major genetic resource to employ gene expression dosage variations (Alonge et al., 2020). SV includes large sequence deletions/insertions, inversions, duplications, and chromosomal rearrangement. Different from SNPs, these gene-associated SVs located in cis-regulatory regions always cause expression dosage changes of corresponding genes and further produce genetic and phenotypic changes. SV was recently reported to be involved in the formation of many traits in plants and is believed to play an important role in plant evolution, crop domestication and improvement (Alonge et al., 2020; Li et al., 2018; Lye and Purugganan, 2019; Rodriguez-Leal et al., 2017). In our study, a 416-bp sequence deletion—a type of SV—that lies in the cis-regulatory region of *AFF* down-regulated the expression of *AFF* and led to its dosage effect as the all-flesh fruit trait (**Fig. 5 and 6**). As exemplified in this study, SVs present as useful quantitative variants, which might be used in next-generation breeding strategy through genetic engineering in the future (Alonge et al., 2020; Rodriguez-Leal et al., 2017; Swinnen et al., 2016).

The expression variation of *AFF* may also contribute to the fleshy fruit evolution in Solanaceae and provides insights into the fruit type evolution among plants. It was proved by archaeology and molecular biology that fleshy fruit plants evolved from dry fruit plants, but the molecular mechanisms responsible for the shift from dry plants to fleshy fruit plants remain unknown (Kumar and Khurana, 2014; Maheepala et al., 2019; Seymour et al., 2008). Therefore, revealing the genetic basis and mechanism underlying the alteration process between fruit types is critical for understanding the evolution of biodiversity. However, the lack of intermediate or transition fruit types has limited the research progress (Annette et al., 2011; Wang et al., 2015). Comparative genetic analysis has shown that there are widespread genomic synteny and collinearity of genes among *Solanaceae* species, especially in Solanaceae vegetable crops (potato, tomato, capsicum, and eggplant), whose fruits show many similar characteristics. However, eggplant and capsicum do not have the same jelly-like tissue in their locule as that of tomato. Those differences in fruit development could also be caused by gene expression variations, other than functional variations of genes (Kim et al., 2014). For example, there is more locule gel in the wild tomatoes *S. lycopersicum var. cerasiforme* and *S. pimpinellifolium* than in cultivated tomatoes (Lemaire-Chamley et al., 2005). We found that the expression of the *AFF* gene in wild tomatoes is also higher than that in cultivated tomato (Tomato Genome Consortium, 2012). The example suggests a positive relationship between the quantity of liquefied locule tissues and the expression level of the *AFF* gene through the process of tomato domestication and breeding.

To summarize, the all-flesh fruit tomato whose locule tissue changes from a jelly-like substance to a solid-state cavity, was found to be caused by an SV of a 416-bp sequence deletion in the cis-regulatory region of the *AFF* gene. The SV mutation reduced the expression dosage of *AFF*, which further impacted the normal liquefication process of locule tissue through the altered expression of subsequent key genes and the subsequent changes in the metabolic components of tomato. Our findings are valuable for revealing the mechanism that underlies changes inside tomato fruit and also shed new light onto the evolution of berry fruit plants. In the future, with systematic studies on the dosage effects of *AFF* expression, accompanied by comparative genomic analysis between plant species of different fruit types and extensive research on the formation and development processes of fruit locule tissues, the evolutionary mechanism of nightshade family fruits and even the berry fruits of different plants will be revealed in depth.

## MATERIALS AND METHODS

### Plant Materials, Growth Conditions, and Phenotyping

Tomato (*S*.*lycopersicum*) plants were cultivated at the Institute of Vegetables and Flowers, Chinese Academy of Agricultural Sciences (IVF-CAAS), Beijing, China, during the natural growing season and under greenhouse conditions. Seeds of *S. lycopersicum* cv. 06-790 and 09-1225 (all-flesh cultivar) were from our own stocks. Seeds of *S. lycopersicum* cv. LA4069, H1706 (LA4345, Heinz 1706-BG; this line was used for the tomato genome sequencing project) and Mico-tom (LA3911) were obtained from the Charles M. Rick Tomato Genetics Resource Center (TGRC) at the University of California, Davis.

The all-flesh line 06-790 was crossed with the wild type, LA4069, to generate F_1_ progeny, and F_2_ progeny were derived from self-pollination of the F_1_ progeny. The F_1_ progeny were crossed with 06-790 to generate BC_1_P1 progeny and crossed with LA4069 to generate BC_1_P2 progeny. These six populations of two crosses were grown for genetic analysis in the greenhouse in spring 2015.

The BC_2_S_1_ progeny were developed from the all-flesh line 06-790 as donor parents, with continued backcrossing to H1706. When the fruit was ripe, the phenotype and characteristics of locule tissue were investigated (10 fruits per plant). The all-flesh fruit individuals of BA-130 and BA-150 and the WT individuals of BA-124 and BA-128 were selected for qPCR.

The NILs of *aff* were derived from BC_6_S_2_ plants generated by 06-790 continual backcrossing to H1706. Among the NILs, BA-1 and BA-2 are *aff* lines and BA-4 and BA-6 are normal lines, and these were used for seed germination. BA-1 and H1706 were also used for morphology research as well as transcriptome and metabolome profiling. Data were analyzed with Excel 2010.

### Paraffin Sectioning and Transmission Electron Microscopy

The locule tissue was cut into 1 mm×2 mm cuboids and fixed in FAA (5% acetic acid, 5% formaldehyde, 50% ethanol, 5% glycerin mixture) for 24 h at room temperature. After dehydration, embedding, slicing, and pretreatment, sections were dyed using safranine and fast green double dye. The paraffin sections were visualized and photographed with and OLYMPUS IX71 microscope.

### Genome Sequencing, SNP, and SV Calling

For rapid identification of the mutation conferring all-flesh fruit in 06-790, we used MutMap, a method based on whole-genome resequencing of bulked DNA of F_2_ segregants (Takagi et al., 2013). We designed two mixed DNA pools that combined the 30 F_2_ progeny that had the all-flesh phenotype and normal phenotype. The DNA pools were subjected to whole-genome resequencing using an Illumina GAIIx DNA sequencer at Beijing Berry Genomics Co., Ltd. The sequencing depth was approximately 20-fold coverage for the two parental lines and approximately 30-fold coverage for the two mixed DNA pools. The paired-end reads of 06-790, LA4069, and the mixed DNA pools were mapped to the tomato reference genome (SL4.0 build; Tomato Genome Consortium, 2012) using Burrows-Wheeler Aligner version 0.7.10-r789 with default parameters (Li and Durbin, 2009).

### Promoter sequence conservation of *AFF* orthologous genes

Syntenic orthologous genes of *AFF* among Solanaceae crop species, *S. lycopersicum, S. pennellii, S. tuberosum, S. melongena*, and *C. annuum*, were determined by the SynOrths tool (Cheng et al., 2012); they are *Sopen06g023350, Sotub06g020180, Sme2*.*5_02049*.*1_g00007*.*1*, and *Capang01g002169*. Then, 5-kb upstream sequences (promoter region) of each of the five orthologous genes were extracted from the genomes of the five species. These sequences were further aligned by MUSCLE (Edgar, 2004). Aligned sequences were further submitted to calculate the conservation level of each aligned nucleotide and then averaged by a 50-bp sliding window with a step of 10 bp. The average values are plotted in **Figure 4a** to show the conservation level of the local regions of the promoter sequences among *Solanaceae* crop genomes.

We further investigated the sequence conservation of the *AFF* gene in the tomato population using the published variome datasets of 360 tomato samples (Lin et al., 2014). We calculated the π values for the 5’UTR, gene body, and 3’UTR for all 35,768 tomato genes in the genome of *S. lycopersicum* with the variome datasets. The distributions of the π values in the three regions (5’UTR, gene body, and 3’UTR) were further plotted as bean-plots with the R package. The locations of the π values of the *AFF* gene were then clearly observed in the background of all tomato genes (**Figure 4b**).

### RNA Extraction and Quantitative Polymerase Chain Reaction (qPCR)

With the use of specific primers and probes, different tomato lines were detected by real-time PCR. They included all-flesh tomato lines BA-130, BA-150, 06-790, and 09-1225, normal tomato lines BA-124, BA-128, LA4069 and H1706, and F_1_ progeny from the crossing of 06-790 and H1706. They were grown in the greenhouse in autumn 2017, and RNA was collected from locule tissues at 7 DAF, 10 DAF, 15 DAF and 25 DAF. The internal reference gene was *SIFRG27* in tomato, the locus was *Solyc06g007510*, and the primer sequence was F (5’-3’): CTCTCTGTTGACAGACCCA; R (5’-3’): GAGTCCAGCTACGAGCAGTG (Cheng et al., 2017).

The primer sequence of *AFF* was F (5’-3’): GCATCTGGTTGGTGAAGG; R (5’-3’): ATCTGATTCTGCTGATGCC. The primers were designed by Roche LCPDS2 software and were synthesized by Beijing TsingKe Biological Technology Co., Ltd. cDNA was obtained from total RNA by using Prime script RT, the reverse transcription kit of Takara Bio Inc. The qRT-PCR was completed on an ABI Prism®7900 qRT-PCR operating system of Applied Biosystems, according to the instructions of the SYBR Prime Script RT-PCR kit. The qRT-PCR and 2^-ΔΔCt^ method were used to analyze the expression of the selected gene.

### Relative Activity of Luciferase

To confirm the function of the 416-bp deletion in promoting the expression of associated geme, the dual luciferase reporter gene assay was used to check the expression difference. Based on the PAS (PCR-based accurate synthesis) method, full-length splicing primers were designed, and the protective base synthesis gene promoters (Del and WT), designed at both ends of the primers, were inserted in sites between PvuII and KpnI in plasmid pGreenII 0800-luc. The recombinant plasmid pGreenII 0800-luc-promoter (Del) was transferred into the epi400 clone strain, and the recombinant plasmid pGreenII 0800-luc-promoter (WT) was transferred to the Top10 clone strain. The sequence of the recombinant plasmid was verified by the sequence of the positive clones.

Monoclones were selected for PCR verification after plasmid transformation. *Nicotiana benthamiana* leaves (one month old) were transiently infected by positive strains using an Agrobacterium-mediated method. Each group was set with three replicates. The activity of the dual luciferase reporter gene was detected after three days. The transcriptional regulation was determined by the activity ratio of firefly luciferase and Ranilla luciferase, that is, the relative activity of luciferase.

### Transcriptome and Metabolome Profiling

Metabolome profiling was carried out using a widely targeted metabolome method by Wuhan Metware Biotechnology Co., Ltd. (Wuhan, China) (http://www.metware.cn/). Briefly, the tomato tissues were lyophilized and ground into fine powder using a mixer mill (MM 400, Retsch) with a zirconia bead for 1.5 min at 30 Hz. Then, 100 mg tissue powder was weighed and extracted overnight with 1.0 mL 70% aqueous methanol at 4°C, followed by centrifugation for 10 min at 10,000 g. All supernatants were collected and filtered with a membrane (SCAA-104, 0.22 mm pore size; ANPEL, Shanghai, China, http://www.anpel.com.cn/) before LC-MS analysis. Quantification of metabolites was carried out using a scheduled multiple reaction monitoring method (Wei et al., 2013; Zhu et al., 2018). In the data analysis process, unsupervised PCA (principal component analysis) was performed by function prcomp within R (version 3.5.0, www.r-project.org). The data were unit variance scaled before performing unsupervised PCA. The HCA (hierarchical cluster analysis) results of samples and metabolites were presented as heatmaps with dendrograms, while Pearson correlation coefficients (PCC) between samples were calculated by the cor function in R. Both HCA and PCC were carried out by R package pheatmap (version 1.0.12). Identified metabolites were annotated using KEGG compound database (http://www.kegg.jp/kegg/compound/); annotated metabolites were then mapped to the KEGG pathway database (http://www.kegg.jp/kegg/pathway.html). Pathways with significantly regulated metabolites were then fed into MSEA (metabolite set enrichment analysis).

For the RNA-seq experiments, a total amount of 3 µg RNA per sample was used as input material for the RNA sample preparations. Sequencing libraries were generated using the NEBNext^®^ Ultra™ RNA Library Prep Kit for Illumina^®^ (NEB, USA) following the manufacturer’s recommendations, and index codes were added to attribute sequences to each sample. The constructed libraries were then sequenced on an Illumina Hiseq platform, and 125 bp/150 bp paired-end reads were generated. Transcriptome profiling was performed as described previously (Ying et al., 2020). Briefly, clean reads were obtained using a Hiseq-X-ten sequencing platform, mapped to the tomato reference genome (Version 4.0) using Hisat 2 (Daehwan et al., 2015), and then normalized to TPM (tags per million reads) reads by StringTie (Pertea et al., 2015). Samples under different combinations were analyzed by MeV (Version 4.9) with the *k*-means method (Gasch and Eisen, 2002). The normalized expression values of genes and metabolites were calculated by dividing their expression level at different time points/tissues. Hierarchical clustering (HCL) and principal component analysis (PCA) were performed to facilitate graphical interpretation of relatedness among different time points/tissue samples. The transformed and normalized gene and metabolite expression values with z-scores were used for HCL and PCA. We used the Pearson’s correlation algorithm method (Bishara and Hittner, 2012) to construct a transcription factor-related gene and metabolite regulatory network. Mutual information was used for calculating the expression similarity between the expression levels of transcription factors and genes, and metabolite pairs were calculated by R software. All the associations among transcription factors, genes, and metabolites were analyzed by Cytoscape software (Kohl et al., 2011).

## Author Contributions

J.L. and L.L. designed and organized the study. L.L., J.B., J.L., X.L., J.H., C.P., S.H., J.Y., and M.Z. conducted the research. F.C., K.Z., L.L., and Y.Z. analyzed the data. All authors discussed and interpreted the results. L.L., F.C., and J.L. wrote the paper. L.L. agrees to serve as the author responsible for contact and ensures communication.

## Acknowledgements

We thank Zhenxian Zhang and Wencai Yang (both China Agricultural University), and Jianchang Gao, Zhognhua Zhang and Xiaowu Wang (all Institute of Vegetables and Flowers, Chinese Academy of Agricultural Sciences) for critical comments.

